# Cortico-Hippocampal phase–amplitude coupling is a signature of learned audiovisual associations in humans

**DOI:** 10.64898/2026.01.20.700563

**Authors:** Arthur Borderie, Laurence Martineau, Paule Lessard Bonaventure, Corentin Labelle, Philippe Albouy

## Abstract

Phase-amplitude coupling (PAC) has been proposed as a neural mechanism for representing sequential information in memory, binding memories, coordinating activity across distributed brain regions, and parsing complex sensory inputs. Importantly, the frequencies contributing to PAC can vary flexibly across brain regions and cognitive states. Here, we asked whether the pattern of coupling across these dynamically varying frequencies can carry information about learned representations. To address this question, we used human intracranial recordings acquired before, during, and after an audiovisual associative learning task. We first identified the emergence of task-related theta- and gamma-band activity within cortico-hippocampal networks during learning and showed that theta-gamma PAC strength within these networks increased following learning. Importantly, PAC comodulograms contained association-specific information: learned associations could be decoded across hippocampal, temporal, and frontal networks during and after learning, with significantly greater decoding accuracy than that obtained from cue-only PAC before learning and from models trained on theta and gamma magnitude alone. This dissociation suggests that the associative information encoded in PAC comodulograms was not reducible to cue-evoked activity or to either of its constituent frequency bands taken independently. Finally, decoding accuracy of models trained on PAC post-learning was positively correlated with behavioral performance. Together, these findings suggest that PAC contains content-specific information in the human brain and identify it as an endogenous neural signature of audiovisual associative learning.

## Introduction

Neural oscillations organize brain activity across temporal and spatial scales, providing windows into how the brain can encode, store, and exchange information[1, 2]. A central hypothesis in systems neuroscience is that cross-frequency coupling (CFC)[3, 4], and more specifically Phase-Amplitude Coupling (PAC)[4, 5], constitutes a computational mechanism enabling information coding in the human brain. PAC refers to the statistical dependency between the amplitude of high-frequency oscillations and the phase of slow oscillations, a dynamic thought to coordinate neural ensembles across different temporal windows[6]. Over the last decade, PAC has been implicated in a broad range of cognitive functions. Lisman & Idiart[7] proposed a model in which the coupling between gamma bursts and the phase of theta oscillations allows multiple items to be represented sequentially within a single theta cycle, a mechanism supported by later human intracranial studies demonstrating theta-gamma interactions[8–12] during working memory maintenance[8, 11, 13]. PAC has also been proposed as a mechanism for selective inter-areal coupling[3–5, 14], enabling the coordination of distant cortical regions, with high-frequency activity modulated by both local and distant low-frequency phase. For sequential episodic memory formation, PAC has been proposed to organize the temporal structure of memory traces[15–18] in both rodent and human studies[15, 18, 19]. PAC has also been proposed as a mechanism for associative binding, whereby the coupling between slow oscillatory phase and high-frequency activity coordinates the simultaneous representation of distinct items across neural populations, enabling the formation of item-item associations that underlie episodic memory[19]. Finally, PAC has been linked to parsing temporally complex sensory inputs such as speech[3, 20, 21], where the phase of low-frequency oscillations aligns with syllabic or phonemic structure to encode fine-grained speech features[3, 21, 22]. Importantly, PAC can arise across a broad and dynamic range of phase- and amplitude-giving frequencies[8] that vary as a function of cognitive state and task demands. Moreover, the relationship between low-frequency phase and gamma amplitude is not fixed across brain regions. For example, recordings in bats have shown stronger delta-theta/low-gamma coupling in auditory cortex, whereas frontal-auditory cortex exhibits preferential coupling between delta phase and high-gamma amplitude[23]. Thus, rather than reflecting a fixed frequency relationship, PAC may provide a flexible representation that is shaped by the underlying brain region and cognitive demands. This flexibility provides a potentially rich representational space in which distinct patterns of cross-frequency coupling could reflect different aspects of the information being processed. PAC should therefore not be viewed solely as a unitary coupling phenomenon captured by a single phase-amplitude pair or by its overall strength, but also as a multidimensional pattern distributed across frequency combinations. To date, however, this rich representational space has not been systematically exploited. Here, we ask whether the frequency-specific patterns of PAC contain information about the content being represented, and whether this information emerges with learning and relates to behavior. For PAC to serve as a genuine information-coding mechanism, its patterns should meet several criteria: (i) item specificity, learned associations should elicit distinct PAC features in relevant brain networks; (ii) these PAC patterns should thus be decodable, allowing classifiers to identify learned-association identity based solely on PAC features (comodulogram); (iii) behavioral relevance. To our knowledge, these criteria have not been tested systematically to date. In the present study, we address this gap by investigating whether PAC patterns in cortico-hippocampal networks carry association-specific (during and after learning) information in humans using an audio-visual associative learning task. Using human intracranial recordings acquired before, during, and after an audiovisual associative learning task, we show that theta-gamma PAC comodulograms within cortico-hippocampal networks contain association-specific information that can be decoded during and after learning, exceeds the information available from cue-only activity or from either constituent frequency band alone, and correlates with behavioral performance. These findings suggest that PAC carries content-specific information in the human brain and identify it as an endogenous neural signature of audiovisual associative memory formation.

## Results

SEEG recordings were obtained from 17 neurosurgical patients (Fig. 1A) with focal drug-resistant epilepsy as they performed an audio-visual associative learning paradigm (Fig. 1B). Participants were initially exposed to auditory (Sound Exposure, SE) and visual (Picture Exposure, PE) stimuli before learning (passive perception). They then learned five arbitrary associations (Associative Learning, AL) between these sounds and letters, with each sound and letter presented simultaneously. AL was followed by two test conditions: during Phase 1 we presented auditory cues and the participants had to press the corresponding letter on the keyboard; During Phase 2, we presented a visual cue and the participants were required to generate mentally the corresponding sound. Then a sound was presented and the participants had to decide if the sound matched or mismatched with the sound that had mentally generated (see Methods for details).

**Fig. 1.**
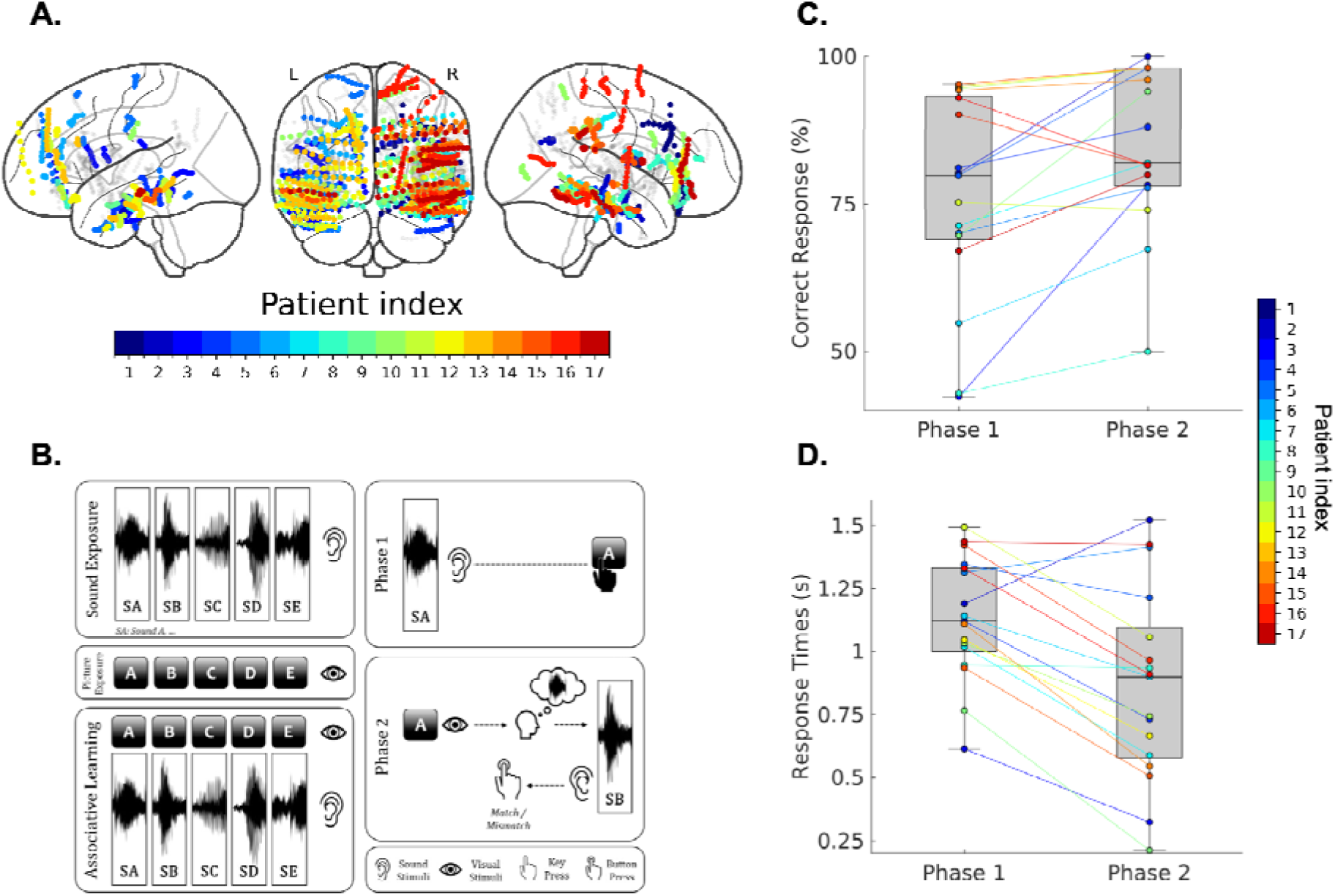
Implantation schema, task and conditions, task performance. A. Implantation schema of all participants represented in the MNI space (N = 17). Colored circles represent SEEG contacts of individual participants (one color per participant). B. The task consisted of five different conditions: Sound Exposure (SE) corresponding to the passive listening of five sounds presented sequentially (by blocks of 5 repetitions); Picture Exposure (PE) corresponding to the passive visual perception of five letters displayed sequentially on the screen (by blocks of 5 repetitions); Associative Learning (AL), sound/letter pairs were presented simultaneously (and different pairs presented sequentially by blocks of 5 repetitions) and participants were required to learn these associations; Phase 1, auditory cues were presented and the participants had to press the corresponding letter on the keyboard and; Phase 2, After a visual cue, participants were required to imagine the corresponding sound and then decided whether a presented sound matched their mental prediction. . C. Accuracy (% correct responses) for Phase 1 and Phase 2 (N = 17, 17). Colored circles represent different participants (one color per participant). D. Response time (s) for Phase 1 and Phase 2 (N = 17, 17). Colored circles represent participants (one color per participant). Error bars reach the maximum et minimum values.

Task performance was evaluated by both the percentage of correct responses and response times (s) for Phase 1 and Phase 2 (Fig. 1C and 1D). To determine whether participants in Phase 1 and Phase 2 performed above chance level (chance at 20% for Phase 1, 50% for Phase 2), two one-sample *t*-test were computed. For both Phase 1 and Phase 2 the percentage of correct responses were significantly above chance with respectively *t*(16) = 13.393, *p* < 0.001 for Phase 1 and *t*(16) = 10.634, *p* < 0.001 for Phase 2.

## Spectral fingerprints of associative learning

To minimize potential confounds from stimulus-locked responses in both time-frequency and PAC analyses, we removed the evoked response from each trial by subtracting the task specific trial-averaged waveform. This step was performed to ensure that oscillatory analyses primarily captured induced activity rather than phase-locked evoked responses[8]. We then investigated time-frequency activity for the associative learning task (see Methods) to establish the presence and temporal evolution of task-related frequency-specific activity during learning. This was done for SEEG contacts located in a priori regions of interest where robust task-relevant activity can be expected[11]: these corresponded to SEEG contacts located in the bilateral superior temporal gyri (Fig. 2A; N_participants_= 14; N_contacts_= 77), middle occipital gyri (Fig. 2B; N_participants_ = 3; N_contacts_ = 5), and hippocampus (Fig. 2C; N_participants_ = 16; N_contacts_= 43), according to the AAL3 atlas. For these regions of interest, the emergence of oscillatory patterns was evaluated across channels by first averaging trials within each channel and participant, then comparing post-stimulus activity to the -300 to 0 ms pre-stimulus baseline across participants, using t-tests with false discovery rate (FDR) correction across time and frequency. In both the superior temporal and middle occipital regions, we observed significant gamma activity (30-150 Hz) beginning approximately 50 ms after stimulus onset. In bilateral hippocampi, significant sustained theta/alpha activity (4-12 Hz) emerged following stimulus onset and lasts around 400 ms. These results demonstrate the presence and temporal evolution of frequency-specific activity in regions where robust task-related activity was expected, thereby supporting the use of these frequency components in the subsequent PAC analysis performed across the entire network. These time-frequency analyses, performed for the associative learning task, established the presence and temporal evolution of task-related activity in the theta (in bilateral hippocampi) and gamma (in sensory regions) frequency ranges in task related regions of interest. This provided an empirical basis for defining the frequencies used for the phase- and amplitude-giving components of the PAC analysis.

**Figure 2:**
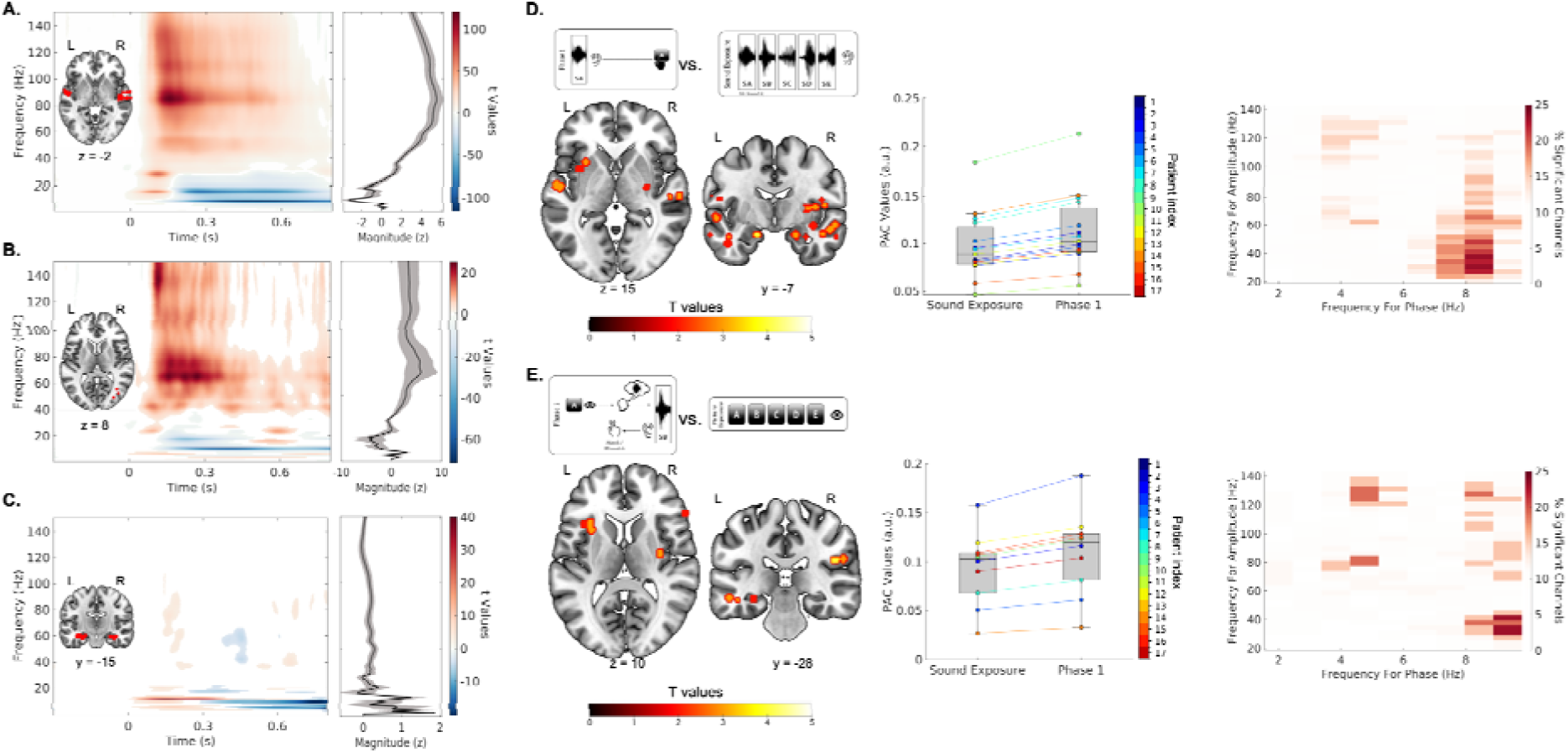
Spectral fingerprints of associative learning. A. Left Panel: T-values in the time-frequency domain (t-tests relative to the -300 to 0 ms pre-stimulus baseline, FDR corrected across time and frequency), computed across participants with SEEG contacts located in the superior temporal gyrus (after averaging trials within each contact), displayed on the single-subject T1 in MNI space provided by SPM12) over the trial time window -300 to 800 ms for the AL condition (N_participant_ = 14; N_contact_ = 77). Right panel shows the PSD, power spectral density (z-score) averaged over a trial time window (50 to 250 ms). Grey shading indicates the standard error of the mean (s.e.m.). B. Same as A. for SEEG contacts located in the middle occipital gyrus (N_participant_ = 3; N_contact_ = 5). C. Same as A for SEEG contacts located the Hippocampus (N_participant_ = 16; N_contact_ = 43). D. Left panel, SEEG contacts displayed on an MR volume (MNI-152), with 4-mm-radius spheres representing spatial clusters of significant contacts showing increased PAC strength across subjects and electrodes for Phase 1 vs. SE contrats (N_participant_ = 15; N_contact_ = 95). Middle panel, Illustration of averaged PAC value per subject for Phase 1 and SE for significant SEEG contacts displayed in A. (colored dots illustrate individual participant). Error bars reach the maximum and minimum values. Right Panel: Illustration of the % of significant electrodes showing an increase post learning vs. pre learning in the comodulogram space E. same as D. for Phase 2 vs. PE (N_participant_ = 10; N_contact_ = 26).

We next asked whether the strength of coupling between these oscillatory components could be specifically modulated after vs. before learning. Phase–amplitude coupling (PAC) was estimated using the direct PAC metric implemented in Brainstorm, based on the mean vector length approach[5, 24] as normalized by Özkurt and Schnitzler (2011) ^24^. Low-frequency phase (2–10 Hz) and high-frequency amplitude (20–150 Hz) were extracted using frequency-domain Gaussian bandpass filters (chirplet filters), retaining only frequency pairs where the high-frequency component exceeded twice the low-frequency component. PAC was quantified over a single 0–600 ms post-stimulus window, yielding one value per frequency pair (comodulogram), contact, and trial. To identify SEEG contacts showing increased PAC after learning, we contrasted post-learning conditions (Phase 1 and Phase 2) against pre-learning conditions (SE and PE) using paired t-tests (p < 0.05, two-tailed) across the comodulogram space, implemented in FieldTrip. Importantly, the pre- and post-learning conditions involved the same sensory stimuli, such that the critical difference between them was the acquisition of the associative relationships post learning. For each SEEG contact and participant, we contrasted Phase 1 (post-learning, involving an auditory stimulus) with SE (cue-only before learning, involving an auditory stimulus) and Phase 2 (post-learning, involving a visual stimulus) with PE (cue-only before learning, involving a visual stimulus) to isolate learning-related changes in PAC. These contrasts were computed for each phase-amplitude frequency coordinates of the comodulogram and performed independently for each contact using cluster-based permutation testing (1,000 permutations). This procedure yielded, for each participant, a set of SEEG contacts showing increased PAC after versus before learning. To evaluate the robustness and generalizability of these effects across participants, we subsequently applied a spatial cluster-based approach in the MNI space, considering only SEEG contacts that overlapped spatially across a minimum of two different electrodes or participants (see Methods and[11]). For visualization PAC values at significant phase-amplitude frequency pairs were pooled across the significant contacts and displayed as box plots to illustrate the magnitude and distribution of the observed effects (Fig. 2D and 2E). In parallel, the corresponding comodulograms displayed on the right panel of Figure 2D and 2E show the percentage of significant contacts at each phase-amplitude frequency pair to provide a descriptive visualization of the frequency distribution of the effects in the comodulogram space. Overall, this analysis revealed a significant increase in PAC strength following associative learning during Phase 1 relative to SE in 95 contacts across 15 participants (Fig. 2D), with significant SEEG contacts clustered in the middle and inferior temporal gyri, the left cingulate gyrus, and the right posterior hippocampus. These included right STSda, PoI2, A1, TE1a, TE2a, PH, H, and IFSp, along with left TE1m, 9a, AAIC, and A4 according to the Glasser’s Atlas[25]. PAC strength also increased during Phase 2 relative to PE in 26 contacts across 10 participants (Fig. 2E), with effects located in the left temporal gyri, extending from lateral to medial and parahippocampal regions, and in the left anterior dorsal insula, as well as the right anterior hippocampus, right inferior frontal gyrus, right superior temporal gyrus, and right posterior insula. These comprised left H, PHA2, MI, and FOP4, and right H, PoI1, A5, IFSa, PF, MI, FOP4, and Ig according to the Glasser atlas.

### PAC carries association-specific information in cortico-hippocampal regions during and after learning

Having confirmed that PAC strength is modulated after associative learning, we next asked whether PAC patterns (i.e. the comodulograms themselves) carry association-specific information during and after learning. PAC was estimated using the direct PAC metric[5, 24] implemented in Brainstorm[26], over a 0-600 ms post-stimulus onset window, yielding one PAC value per frequency pair per contact per trial (9 phase-frequency × 36 amplitude-frequency bins, 324 features total). For each contact and condition, a multiclass ridge classifier was trained to discriminate among the five learned audiovisual associations (225, 160, and 102 trials for AL, Phase 1, and Phase 2 respectively), using an 80/20 development/held-out test split with five-fold cross-validation for hyperparameter selection within the development set. Classification accuracy was evaluated on the held-out test set and assessed against a 1,000-label-permutation null distribution, with the complete classification procedure (including five-fold cross-validation for hyperparameter selection) repeated for each permutation. Contacts exceeding the 95th percentile of this null distribution were considered above chance. As in the univariate analyses, significant contacts were projected onto MNI space and subjected to the same spatial clustering procedure to identify networks showing spatially reproducible decoding across participants (Fig. 3; see Methods for details).

**Fig. 3.**
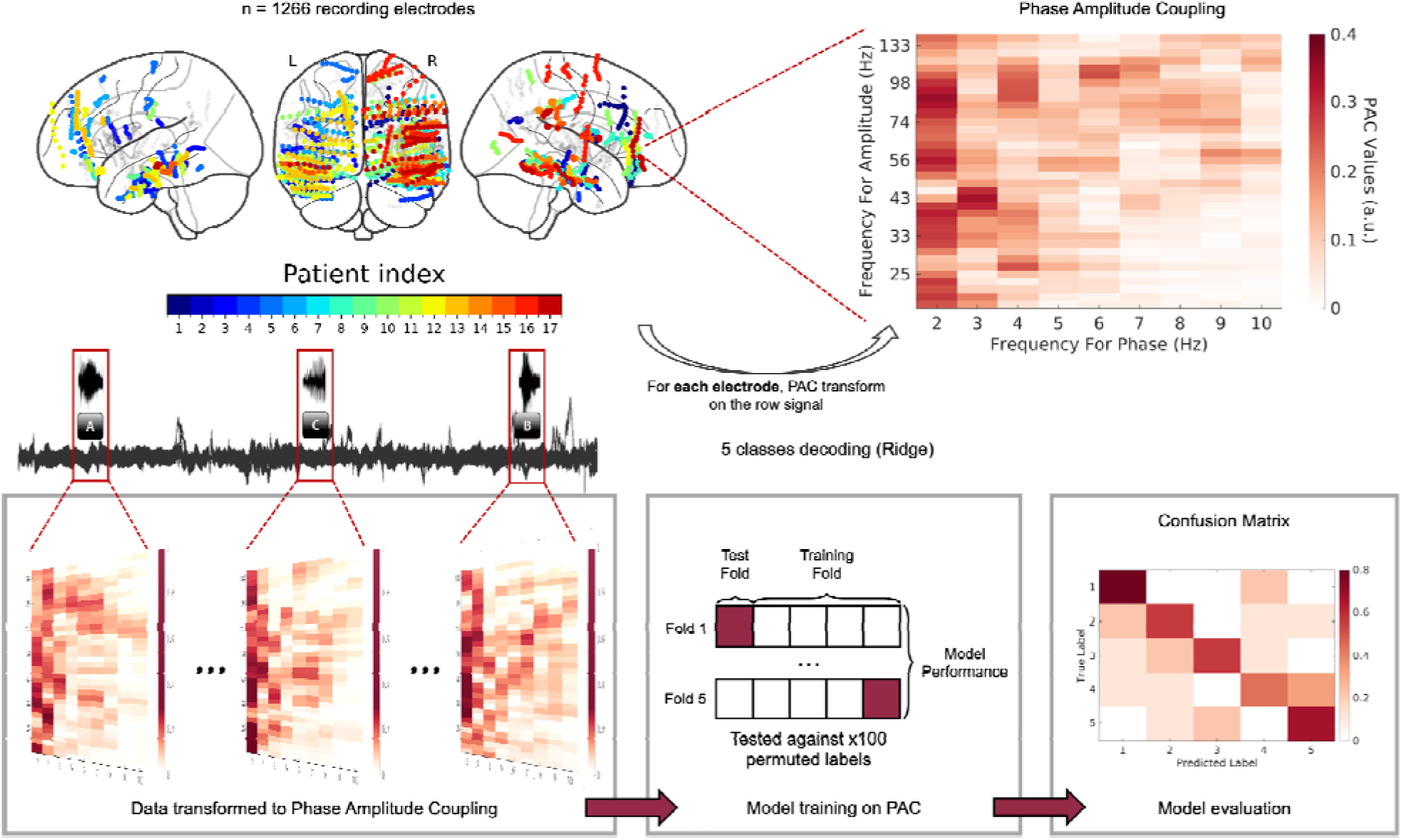
Illustration of the PAC-based multiclass decoding pipeline. For each SEEG contact (top left, color-coded by participant), the raw signal was transformed into a phase-amplitude coupling (PAC) comodulogram (top right), yielding one PAC value per low/high-frequency pair (9 × 36 = 324 features) over a 0-600 ms post-stimulus window. One comodulogram was computed per trial, resulting in a set of comodulograms stacked across trials for each contact (bottom left). A multiclass ridge classifier was then trained to discriminate among the five learned audiovisual associations, using an 80/20 development/held-out test split with five-fold cross-validation for hyperparameter selection within the development set (bottom center). Classification performance was evaluated on the held-out test set and assessed against a null distribution of 1,000 label-permutation accuracies, with the full procedure repeated for each permutation. Decoding performance is summarized by a confusion matrix (bottom right), where diagonal entries reflect correct classification of each association. Contacts exceeding the 95th percentile of the permutation null distribution were considered above chance, and spatial clustering across participants was applied to identify networks with spatially reproducible decoding.

During the associative learning condition (AL; Fig. 4A), mean classification accuracy was 0.337 ± 0.087% for 6 contacts in 6 participants, with significant classification decoding in bilateral hippocampi (H, PHA1) and right anterior insula (MI). For the post-learning conditions, classification for Phase 1 (Fig. 4B), mean accuracy was x 0.302 ± 0.079% across for 24 contacts in 13 participants, with significant SEEG contacts clustered in the left inferior frontal cortex (a47r), inferior temporal gyrus (TE1a) and hippocampus (H) and in the right superior temporal gyrus (A5, A1, STSva), posterior insula (TPOJ1), middle insula (MI, PoI2) and para hippocampal area (PHA3). For Phase 2 (Fig. 4C), mean accuracy was 0.343 ± 0.101% across 14 SEEG contacts in 8 participants, with SEEG contacts clustered in the left ventral middle insula (AAIC) and in the right middle temporal gyrus (TE1m, TE1p), inferior frontal cortex (45), posterior cingulate (RSC, 23c) and hippocampus (H). Across channels, the median standard deviation of test accuracy across the five folds was 0.07 (IQR: 0.059–0.098).

**Figure 4:**
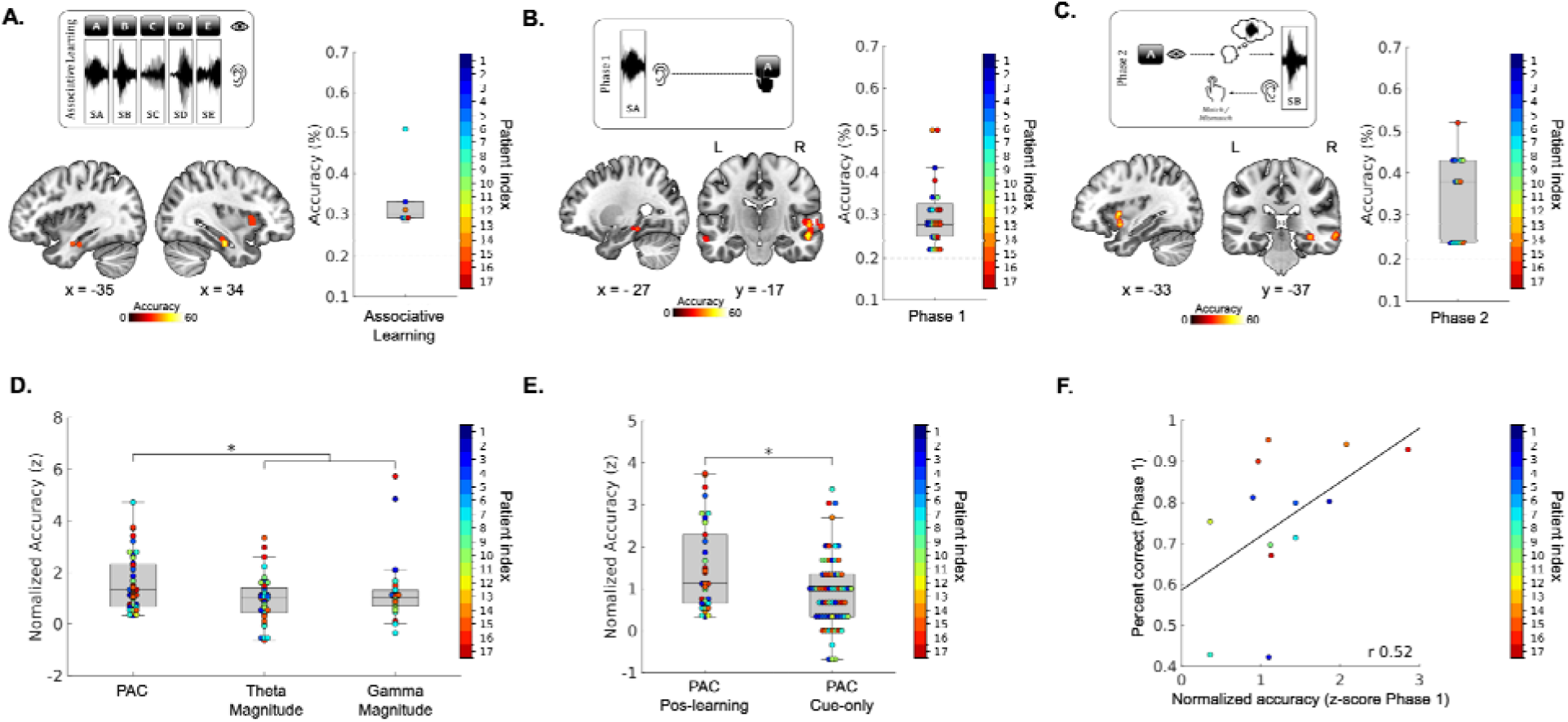
PAC decoding results. A. Left panel, SEEG contacts displayed on an MR volume (MNI-152), with 4-mm-radius spheres representing spatial clusters of SEEG contacts showing significant decoding accuracy (permutation test, 1,000 permutations) for the AL condition (N_participant_ = 6; N_contact_ = 6). Right panel, illustration of the average decoding accuracy (%) per subject for significant SEEG contacts displayed in A. Colored circles represent individual participants (one color per participant, average decoding accuracy = 0.337; std = 0.087). B. Same as A. for the Phase 1 condition (N_participant_ = 13; N_contact_ = 24, average decoding accuracy= 0.302; std = 0.079). C. Same as A. for the Phase 2 condition (N_participant_ = 8; N_contact_ = 14, average decoding accuracy = 0.343; std = 0.101). D. Normalized decoding accuracy (z-scores, calculated by subtracting the mean of the permutation distribution from the observed accuracy and dividing by its standard deviation) for PAC, theta, and gamma features collapsed across conditions (AL, Phase 1, and Phase 2; 14 participants, 44 contacts). Colored circles represent individual participants (one color per participant). E. Normalized decoding accuracy (z-scores, calculated by subtracting the mean of the permutation distribution from the observed accuracy and dividing by its standard deviation) for PAC post learning (phase 1, Phase 2), and PAC cue only (SE, PE) features collapsed across conditions. Colored circles represent individual participants (one color per participant). F. Scatter plot showing the linear relationship between behavioural performance in Phase 1 (%) and mean PAC decoding accuracy across significant SEEG contacts for each participant (N_participants_ = 13; N_contact_ = 24; Pearson’s r = 0.516, one-tailed p = 0.03). Colored circles represent individual participants

### PAC decoding exceeds single-band alternatives, increases after learning, and correlates with behavioral performance

We then compared decoding accuracy with PAC as features with decoding based on theta or gamma activity alone. Using the same classification framework as for PAC decoding (identical classifier, 5-fold cross-validation, time window, and trials per class; see Methods), classifiers were trained separately on time-frequency representations (Morlet) for theta (4-10 Hz), low-gamma (40-90 Hz), and high-gamma (90-150 Hz) activity, resized to match the dimensions of the PAC comodulograms. Because low- and high-gamma were correlated across contacts in our data (r = 0.52, p < .05), we averaged their decoding scores for each contact into a single gamma score, so that PAC was compared against one theta and one gamma alternative. These analyses were performed on the same SEEG contacts that showed significant PAC decoding (above chance, 1000 permutation test) and after spatial clustering (44 contacts, 14 participants; Fig. 4A, 4B and 4C), separately for each condition. Decoding accuracy for each feature was normalized against a permutation-based null distribution (1,000 permutations) by converting observed accuracy to a z-score, quantifying how far performance deviated from chance in units of the null distribution’s standard deviation. We then compared the three features (PAC, theta-, gamma-magnitude), while accounting for the non-independence of contacts within participants, with a generalized estimating equation (GEE, participants as clusters, exchangeable working correlation, robust sandwich standard errors), with feature and condition (associative learning, Phase 1, Phase 2) as factors. There was a main effect of feature (sequential Wald χ²(2) = 11.79, p = .003). PAC normalized decoding accuracy was significantly higher than both theta-based decoding (+0.66 z, 95% CI [0.29, 1.04], z p = .017) and gamma-based decoding (+0.60 z, 95% CI [0.23, 0.98], z p = .026), whereas theta and gamma did not differ from each other (p = .68; Fig. 5A). Because only 14 participants contributed, we also applied Kauermann-Carroll small-sample correction, which widened the intervals but preserved the direction of effects (PAC vs. theta: 95% CI [0.14, 1.19], t(5) p = .023; PAC vs. gamma: 95% CI [0.07, 1.14], t(5) p = .034). The feature × condition interaction was significant by the sequential Wald test (χ²(4) = 20.97, p < .001) but did not survive Kauermann-Carroll correction (F(4, 5) = 4.59, p = .063; Fig. 4D).

To test whether decoding after learning exceeds decoding before learning, we compared the same single cue across learning: sounds during passive exposure (SE, pre-learning) with sounds during the first evaluation phase (Phase 1, post-learning), and letters during passive exposure (PE, pre-learning) with letters during the second evaluation phase (Phase 2, post-learning). The stimuli are therefore physically identical before and after learning. The associative learning (AL) condition was not used for this comparison because it contains both cues on every trial. Analyses included contacts showing above-chance decoding after learning (visual: 14 contacts; auditory: 24 contacts; 14 participants). To evaluate whether PAC decoding differed between post- and pre-learning periods, we analysed decoding z-scores with a generalized estimating equation (GEE; R package geepack), with participant as the clustering unit, a Gaussian identity link, and an exchangeable working correlation structure. The model included modality (visual, auditory; effect-coded ±0.5), learning (POST vs PRE; effect-coded ±0.5), and their interaction as fixed effects, allowing us to estimate the population-average PRE-to-POST change in decoding accuracy while accounting for the non-independence of contacts within participants. Cluster-robust sandwich standard errors were used for inference, with p-values derived from z-tests. Because only 14 participants contributed, we additionally applied the Kauermann-Carroll small-sample correction to the sandwich covariance matrix, re-evaluating all effects against a t distribution with degrees of freedom equal to the number of clusters minus the number of model parameters (df = 10). PAC decoding was higher after learning than before (β = 0.40 z, 95% CI [0.01, 0.78]; GEE with participant as cluster, robust SEs, z-test p = .02). There was no significant modality × learning interaction (β = 0.30 z, z p = .461), indicating that the PRE-to-POST increase did not differ between visual and auditory contacts. Because only 14 participants contributed, we also applied small-sample corrections, which widened the intervals (Kauermann-Carroll: 95% CI [−0.01, 0.81], t(10) p = .05).

To further evaluate whether PAC patterns contained information related to the strength of the learned associations, we investigated the relationship between post-learning PAC normalized decoding accuracy (see methods) and behavioural performance across participants. Because our hypothesis specifically predicted a positive relationship between the strength of the neural representation and behavioural performance, we used a one-tailed test. In Phase 1, PAC decoding accuracy was positively related to behavioural performance (r = 0.516, one-tailed p = 0.03; Fig. 4F). The same analysis in Phase 2 did not reveal a significant relationship (p > 0.05;). This should be interpreted cautiously because fewer participants had contacts showing significant PAC decoding in Phase 2 (N = 8) than in Phase 1 (N = 13), limiting the power to detect an across-participant association.

## Discussion

We recorded SEEG data from 17 patients with drug-resistant epilepsy before, during and after an audio-visual associative learning task. Our paradigm involved passive exposure to sounds and pictures, associative learning of five sound/letter pairs, and two test phases assessing behavioral recall from auditory cues (Phase 1) and mental sound generation from visual cues (Phase 2). In addition to confirming learning-related modulations of PAC strength, our results demonstrate that PAC in cortico-hippocampal networks fulfills key criteria for a genuine information-coding mechanism. PAC comodulograms carried association-specific signatures that could be decoded in distributed fronto-temporal-hippocampal networks during and after learning, with decoding accuracy exceeding that obtained from either constituent frequency band, theta or gamma, considered independently. PAC decoding was also greater after learning than before, when stimuli were physically identical, but associations had not yet been formed, suggesting that the information carried by PAC comodulograms reflects the learned associative structure rather than sensory properties of the cues alone. Finally, post-learning PAC decoding accuracy was positively correlated with behavioral performance during the first transfer phase, linking the strength of the neural representation to the quality of memory encoding. Together, these findings provide direct evidence that cortico-hippocampal PAC constitutes a content-specific neural code supporting associative learning in the human brain.

### Learning-related enhancements of cortico-hippocampal PAC

We first established the presence of task-relevant frequency-specific activity in a priori regions of interest during associative learning. Significant gamma activity (30-150 Hz) emerged approximately 50 ms after stimulus onset in superior temporal and middle occipital regions, while sustained theta/alpha activity (5-12 Hz) was observed in the hippocampus for approximately 400 ms following stimulus onset. These results confirm the presence and temporal evolution of the frequency components targeted in the subsequent PAC analysis and are consistent with numerous neuroimaging studies showing that the temporo-fronto-hippocampal network integrates sensory representations with mnemonic processes to form, maintain, and retrieve cross-modal associations[11, 27]. Building on this, we then showed that post-learning PAC strength was significantly increased relative to pre-learning, particularly in secondary auditory regions, hippocampal areas and inferior frontal regions. These results are well aligned with numerous neuroimaging studies showing that the temporo-fronto-hippocampal network integrates sensory representations with mnemonic processes to form, maintain, and retrieve cross-modal associations[18, 28–31]. In addition, the observed coupling within hippocampal and parahippocampal regions is consistent with their well-established role in new memory formation, supporting models in which distributed sensory representations are gradually incorporated into hippocampal relational codes to enable flexible associative memory[31, 32]. However, such task-related increases alone do not demonstrate that PAC patterns represent which item, stimulus, or learned association is being processed. To directly test whether PAC serves as a specific representational marker, we applied linear classifiers to PAC features to determine whether item-specific information could be decoded before, during, and after learning.

### PAC transitions from sensory to higher level coding

Multivariate decoding analyses demonstrated a clear spatial progression in the information content carried by PAC. During learning, PAC patterns encoded association-specific information, in bilateral anterior hippocampus and right anterior insula. We interpret this pattern as reflecting the integration of sensory inputs with hippocampal regions, consistent with evidence that hippocampo-cortical theta-gamma interactions support relational memory formation[6, 8–10, 13, 15, 18, 19, 33–35]. Post-learning, we showed that PAC encoded association-specific information, though anatomo-functional network configurations shifted. In Phase 1, significant decoding relied on right secondary auditory regions (A5, A1, STSva), insular, lateral prefrontal areas, and middle/posterior hippocampus. We propose, based on previous studies that this network auditory, prefrontal, and medial temporal region reflect stabilization and retrieval of learned associations[36, 37]. For Phase 2, decoding was observed left anterior insula and right frontal, secondary auditory cortex (TE1m and TE1p), posterior hippocampus and posterior cingulate. This brain pattern could be related to a transformation toward abstract, modality-independent representations of learned associations during recall with potential connection between the posterior cingulate and hippocampus as part of the mnemonic recall process[38]. Consistent with this interpretation, and although not formally tested, the distribution of contacts contributing to significant PAC decoding showed a descriptive shift along the hippocampal long axis across conditions: contacts were predominantly located in the anterior hippocampus during associative learning, and shifted toward more posterior hippocampal regions during Phase 1 and Phase 2 (Figure 4 ABC). This anterior-to-posterior gradient is consistent with the known functional specialization of the hippocampal long axis, whereby the anterior hippocampus supports rapid associative binding during initial encoding and the posterior hippocampus is associated with more precise, stable, and transferable memory representations[39, 40]. This observation, however, warrants formal investigation in future work. Altogether, these results suggest that PAC patterns encode higher-order associative information in distributed cortico-hippocampal-prefrontal networks, supporting theoretical models in which theta-gamma coupling coordinates hierarchical communication to bind stimuli into stable memory representations[6].

### PAC carries memory-related information beyond spectral power and relates to behavioral performance

Within the network of contacts showing significant PAC decoding for AL, Phase 1 and Phase 2, normalized decoding accuracy based on PAC comodulograms exceed that based on theta and gamma magnitude features considered separately. This suggests that PAC may capture associative information that is not fully reflected in either constituent frequency band alone. This is consistent with the view that theta-gamma coupling reflects the active binding of associated items, over and above what can be inferred from broadband power changes. This is also in line with human intracranial studies showing that theta-gamma PAC coordinates mnemonic processing across hippocampal and cortical regions[15, 41]. Note that this analysis was designed as a control to ask whether, within the network of contacts already showing significant PAC decoding, the same associative information could be decoded from theta or gamma magnitude alone. PAC decoding systematically exceeded single-band decoding within these networks, suggesting that the phase-amplitude coupling itself carries information beyond what is available in either constituent frequency band considered independently. However, this comparison should be interpreted with caution, and future work evaluating the relative contribution of PAC and single-band features across the whole brain will be needed to more fully characterize the specificity of this effect. Critically, the superiority of PAC over single-band features cannot by itself rule out the possibility that PAC patterns simply reflect cue-related sensory responses that differ across the five associations. To address this, we compared PAC decoding accuracy for physically identical stimuli before and after learning. The finding that PAC decoding accuracy was higher after learning than before (when the stimuli were physically identical but associations had not yet been formed) provides important evidence that the information carried by PAC comodulograms reflects the learned associative structure rather than the sensory properties of the cues themselves. This learning-related increase was observed across both visual and auditory conditions without a significant modality difference, suggesting that the emergence of association-specific PAC patterns is not driven by modality-specific sensory processing but rather by a supramodal mnemonic representation of the learned pairings. This is consistent with the view that cortico-hippocampal theta-gamma coupling supports the active encoding and consolidation of associative memory traces[18, 28–31]. An additional consideration is that eye movements may influence hippocampal oscillatory dynamics[42], as saccade-triggered phase resetting has been reported in the human hippocampus. Given that the present task likely involved eye movements, future studies combining intracranial recordings with eye tracking could determine whether such oculomotor-related dynamics contribute to the observed PAC patterns.

Finally, we observed a positive relationship between PAC decoding accuracy and behavioral performance during Phase 1, whereas no corresponding association was observed in Phase 2. This effect is consistent with previous reports linking hippocampal PAC to successful episodic encoding, memory updating, and individual differences in memory performance[15, 41, 43]. However, the difference between Phase 1 and Phase 2 should be interpreted cautiously, as the Phase 2 analysis included fewer participants with significant PAC decoding (N = 8 vs. N = 13 in Phase 1), limiting the precision and power of the corresponding analysis. Moreover, the present analyses do not directly test whether the strength of the PAC-behavior relationship differs between retrieval phases. The observed pattern nevertheless suggests that PAC-based representations may be related to memory performance during auditory-cued retrieval, a possibility that should be further evaluated in larger samples with matched behavioral measures across retrieval conditions.

Overall, our findings extend evidence linking PAC to associative binding by showing that coupling patterns discriminate learned associations, rather than reflecting a simple variation of PAC strength with learning or memory success[19, 44]. These observations raise the possibility that PAC reflects not only the engagement of associative processing but also its representational content. However, a binding mechanism and an item-specific representational marker are not equivalent: the former concerns how representations become associated, whereas the latter concerns whether a measured signal distinguishes their content[45]. We propose that item-specific PAC patterns could reflect the recruitment of distinct neuronal assemblies within a shared temporal coordination mechanism. This interpretation is compatible with computational work linking oscillatory coordination to the segregation and selective transmission of representations[46], and the higher PAC decoding accuracy after learning compared to before further suggests that this coupling reflects active mnemonic processing rather than a passive consequence of item-specific sensory responses. The relationship to sequential coding is similarly one of compatibility rather than direct confirmation[47]. Previous findings associate successful sequence memory with the ordering of item-related gamma activity along theta phase[18]. Our results instead identify item-specific differences in the spectral configuration of PAC, without establishing their ordering within an oscillatory cycle, and therefore address a complementary representational dimension (e.g. which item or association is being processed, rather than where it occurs in a sequence). Finally, the distribution of PAC decoding across hippocampal and cortical regions is compatible with hippocampo-cortical accounts of associative representation and retrieval. Human intracranial studies have linked hippocampo-cortical cross-regional PAC to successful encoding[41] and interregional gamma phase synchronization to coordinated hippocampal and cortical reinstatement[48]. Our results complement this distinction by identifying representational information in PAC patterns, without establishing that these patterns mediate interregional transfer or hippocampally driven reinstatement. Thus, PAC’s proposed binding function and its observed representational specificity may be complementary, but demonstrating their mechanistic relationship will require linking item-resolved PAC patterns to interregional coordination and testing their causal contribution to associative memory.

Collectively, our findings demonstrate that PAC patterns in cortico-hippocampal networks constitute a flexible, structured, and potentially behaviorally relevant neural code. Beyond prior work that largely emphasized PAC strength changes, we show that PAC carries representational content, aligns with behavior, and is shaped by individualized and regions specific neural architectures. This positions PAC as a mechanism bridging local cortical computations with distributed cortico-hippocampal networks, supporting learning and memory.

## Materials and Methods

### Resource availability

#### Lead contact

Further information and data that support the findings of this study are available upon request from the corresponding author.

#### Material availability

The raw brain data used in this study (MRI and SEEG recordings) are protected as acquired through clinical assessment and are not available for sharing (CHU de Québec - Université Laval 2022-5890). Data used to generate the figures and MATLAB scripts are available at the following URL (https://osf.io/d8cuy).

#### Data and code availability

Original codes used to process SEEG and behavioral data are publicly available on OSF (https://osf.io/d8cuy), or upon request to the corresponding author.

### Experimental design and populations

#### Patients

SEEG data from 17 neurosurgical patients with drug resistant focal epilepsy (8 females and 9 males, mean age: 31.7 +/− 8.47 years) were collected at the Epilepsy Department of the Enfant Jesus Hospital (Quebec, Canada). The research and clinical teams verbally explained the study protocol, and eligibility was discussed with patients prior to hospitalization. All participants gave verbal assent and signed a written informed consent form. The study adheres to all applicable ethical guidelines, with experimental procedures approved by the Ethics Review Board of the CHU de Québec - Université Laval (2022-5890).

### Method details

#### Behavioral task

Participants were asked to perform an associative audio-visual task, consisting in five distinct conditions: Sound Exposure (SE), Picture Exposure (PE), Associative Learning (AL), Evaluation of Learned Associations 1 (Phase 1), and Evaluation of Learned Associations 2 (Phase 2). In all conditions, stimuli were separated by a 1-s interval followed by a randomly jittered interval of 0.5-1 second. 1) Sound Perception Block (SE): Participants passively listened to five complex environmental sounds (e.g., a door closing, water flowing, etc.; adapted from[49], presented sequentially in blocks of five repetitions (25 trials per sound; sound duration: 500 ms). 2) Picture Perception Block (PE): Participants passively viewed five letters displayed sequentially, presented sequentially in blocks of five repetitions, on a screen (25 trials/letter, letter presentation: 500 ms,). 3) Associative Learning Condition (AL): Participants learned to associate each sound with a specific letter through paired presentations, presented sequentially in blocks of five repetitions (45 trials/sound-letter pair, pair duration: 500 ms). 4) Evaluation of Learned Associations 1 (Phase 1): In each trial, one of the five sounds was played (presented randomly), and participants had 2 seconds to indicate the associated letter using a keyboard (5 predefined keys, 32 trials/sound). 5) Evaluation of Learned Associations 2 (Phase 2): For each trial, a letter was displayed (presented randomly), and participants mentally generated the corresponding sound. After a 1.5-second delay, a sound was presented, and participants judged whether it matched the mentally generated sound (20 trials/category) using a mouse. Stimuli mismatches occurred in 50% of trials, with mismatched sounds selected from the same environmental sound pool (see Figure 2A for trial structure). All auditory stimuli were delivered via headphones, and visual stimuli were presented on a 24-inch monitor with a 60 Hz refresh rate.

#### Procedure

SEEG data were recorded continuously while patients sat approximately 75 cm from the computer screen (1024 × 768 pixels). Before the recording began, each patient received verbal instructions to ensure they understood the task, though no practice trials were conducted. Presentation software (Neurobehavioral Systems, Albany, CA, USA) was used for the delivery of the experimental protocol, to present the auditory and visual stimuli, and to register button presses. Conditions were presented in the same order (SE, PE, AL, Phase 1 and Phase 2) for all participants as some conditions are prerequisite for others. Participants were informed of the block order and if needed were asked to indicate their answers by pressing the correct key on the keyboard (Phase 1) or the correct mouse button (Phase 2). Their responses were recorded during the first 2 s of the inter-trial interval. A feedback was given during the experiment. This feedback was represented as emojis of different emotions and color with green for correct answers, red for wrong answers and blue for misses. Phase 1 and Phase 2 contained an equal number of ‘same’ and ‘different’ trials. For these two conditions, the trials were presented in a pseudo-randomized order; the same trial type (i.e., ‘same’ or ‘different’), could not be repeated more than three times in a row. In the SE, PE, and AL conditions, each item was presented in blocks of five consecutive trials, with blocks cycling through items 1-5. This cycle was repeated five times in SE and PE and nine times in AL.

#### SEEG electrodes and SEEG contacts localization

Patients’ brains were stereotactically implanted with 10 to 18 semi-rigid multi-lead electrodes (AdTech, diameter: 0.8 mm). According to the clinically defined target structure, activity was recorded over SEEG contacts from 2.0 mm wide and 3, 4, 5 or 6 mm apart (depending on the electrode). All participants underwent two 3D anatomical MPRAGE T1-weighted Magnetic Resonance Imaging scan on a a 3T Siemens Trio (Siemens AG, Erlangen, Germany) before implantation and just after the SEEG implantation. The anatomical volume consisted of 160 sagittal slices with 1 mm3 voxel, covering the whole brain. All electrode contacts were identified on the post-implantation MRI showing the electrodes and co-registered on a pre-implantation MRI (ImaGIN toolbox, https://f-tract.eu/software/imagin/). MNI coordinates were computed using the SPM (http://www.fil.ion.ucl.ac.uk/spm/) toolbox. In addition to MNI coordinates, we computed the localization of the SEEG contacts in the Human Connectome Project Atlas[25].

#### SEEG recordings and preprocessing

Intracranial recordings were conducted using a video-SEEG monitoring system (NATUS), which allowed the simultaneous data recording from up to 277 depth EEG electrode sites. The data were bandpass filtered online from 0.1 to 200 Hz and sampled at 512 Hz for all patients. Powerline contamination of the raw data (main 60 Hz, harmonics 120 Hz and 180 Hz) was reduced using notch filtering. Each SEEG signal was then re-referenced to its immediate neighbor through bipolar derivations. Bipolar montage refines local specificity by eliminating signal artifacts common to adjacent contacts (e.g., 60 Hz running through wires and distant physiological artifacts) and cancel effects of distant sources whose signal is spreading to adjacent sites through volume conduction. Using this method, the estimated spatial resolution of each SEEG bipolar contact was of 3-4 mm[50].

#### SEEG signal processing and analyses

##### Signal preprocessing

SEEG data were preprocessed and visually checked by the medical team to reject contacts contaminated by pathological epileptic activity or environmental artifacts. SEEG preprocessing and statistical analyses were performed using Brainstorm[26] (http://neuroimage.usc.edu/brainstorm/) and FieldTrip[51] functions (www.fieldtriptoolbox.org), as well as custom MATLAB codes. Data were epoched to create trials with a window of 500 ms before stimuls onset and 1000 ms after stimulus onset.

##### Evoked response removal

To isolate induced activity from the stimulus-locked evoked response, the task-specific trial-averaged waveform for each participant and SEEG contact was subtracted from the data of each trial prior to subsequent analyses.

##### Time-frequency analysis

Time-frequency analyses based on Morlet wavelets were performed for the associative learning task on SEEG contacts located in a priori regions of interest where robust task-related activity was expected, including bilateral superior temporal gyri, middle occipital gyri, and hippocampi, according to the AAL3 atlas. This analysis was performed to characterize frequency-specific activity in these regions and to establish the frequency bands used for the phase and amplitude components of the subsequent whole-brain PAC analysis. Morlet wavelet transforms were computed separately for each SEEG contact and trial, using a wavelet family defined by f0/σf = 3, with f0 ranging from 1 to 150 Hz in 1-Hz steps. The magnitude of the wavelet transform was then averaged across trials within each contact and averaged across participants, providing an estimate of oscillatory power at each time point and frequency between 1 and 150 Hz. Magnitude was baseline-normalized (z-scored) relative to the −300 to 0 ms period preceding stimulus presentation. Statistical significance was evaluated by performing t-tests against the pre-stimulus baseline (−300 to 0 ms relative to stimulus onset) across participans within three ROI (bilateral superior temporal gyrus, middle occipital gyrus and hippocampi), with FDR correction across time and frequency.

##### Phase-amplitude coupling transformation

Phase-amplitude coupling (PAC) was estimated using the direct PAC metric implemented in Brainstorm, following the normalization approach described by Özkurt and Schnitzler (2011)[24] and inspired by the original mean vector length method of Canolty et al. (2006)[5]. Continuous SEEG signals (sampled at 512 Hz) were first broad-band filtered using an FFT-based band-pass filter with boundaries set to half the minimum nesting frequency and 1.5 times the maximum nested frequency, implemented without phase shift. To reduce edge effects, signals were mirrored before filtering and the mirrored portions removed after filtering. The filtered signals were transformed into the frequency domain via FFT, from which analytic representations were derived by zeroing negative frequencies and doubling positive frequencies. Frequency-specific amplitude and phase were then extracted using frequency-domain Gaussian bandpass filters (implemented as chirplet filters with zero chirp rate in Brainstorm) centered at each frequency of interest, with a fractional bandwidth parameter of 0.15, such that the filter bandwidth scaled proportionally with center frequency. Based on the time-frequency results obtained during learning, low-frequency phase was extracted in the 2-10 Hz range and high-frequency amplitude in the 20-150 Hz range. For each low/high-frequency pair, PAC was quantified as the magnitude of the sum of amplitude-weighted phase vectors, normalized by the L2 norm of the amplitude envelope and the square root of the number of time points, following Özkurt and Schnitzler (2011)[24]. Frequency pairs were retained only when the high-frequency component exceeded twice the low-frequency component. PAC was estimated over a single 0-600 ms post-stimulus onset window, yielding one PAC value per frequency pair per contact per trial, with no temporal binning applied. These values were used directly in subsequent analyses without additional normalization.

##### Phase-amplitude coupling contrasts

To identify SEEG contacts where PAC strength increased post learning conditions (Phase 1 and Phase 2) as compared to pre learning condition (SE and PE) we performed a cluster based contrast in the comodulogram space using FieldTrip function (paired t-tests (*p* < 0.05, two-tailed). To control for multiple comparisons, cluster-based permutation testing (1,000 iterations) was applied at the SEEG contact level, with cluster-forming and cluster-corrected thresholds of *p* < 0.05. To minimize the type I error rate, we applied a spatial clustering across patients.

##### Machine Learning for PAC-Based Neural Decoding

To test whether PAC patterns contained information that differentiated the learned associations, we performed multiclass classification separately for each SEEG contact. For each trial and channel, the PAC comodulogram was represented as a feature vector comprising 324 PAC values (9 phase-frequency bins × 36 amplitude-frequency bins). The five learned associations constituted the five classification categories. with 225 trials during the AL condition, 160 trials during Phase 1, and 102 trials during Phase 2. For each SEEG contact, the data were divided into an 80% development set and a 20% held-out test set, with the class distribution preserved across the two sets. A multiclass ridge classifier was trained using the 324 PAC features. Within the 80% development set, the ridge regularization parameter (α) was optimized over a predefined grid using five-fold cross-validation. The classifier was then refitted on the complete development set using the selected α and evaluated once on the independent held-out test set. Thus, the held-out test data were not used for hyperparameter selection or model fitting. To assess whether classification performance exceeded chance levels, we performed a label-shuffling permutation analysis independently for each channel. Class labels were randomly permuted 1,000 times, and the complete classification procedure was repeated for each permutation, including five-fold cross-validation for selection of α within the development set. Classification accuracy was subsequently evaluated on the corresponding held-out test set, yielding a null distribution of 1,000 accuracy values. The 95th percentile of this permutation-derived null distribution was used as the significance threshold for the observed classification accuracy. Classification performance was considered above chance when the observed accuracy exceeded this threshold. Each SEEG contact was treated as a separate unit of signal analysis and classification was performed separately for each channel.

##### Spatial clustering

Spatial clustering. For both univariate and multivariate analyses, significant contacts from all patients for each task were projected onto a common standardized brain space (MNI-152). A 3D sphere (radius: 4 mm), reflecting the spatial resolution of the bipolar montage, was centered on each significant contact and overlaid onto the MRI volume. Contacts were considered to form a spatial cluster when their 4-mm spheres overlapped. Only clusters comprising at least two overlapping SEEG contacts were retained, whereas isolated significant contacts were excluded to reduce the likelihood that effects driven by a single contact would be interpreted as network-level effects. To ensure that retained clusters reflected genuine cross-participant and cross-electrode convergence, we constrained that each retained cluster included contacts from at least two different participants and from at least two different electrodes. Thus, spatial clustering was used as a group-level criterion to identify spatially reproducible effects across participants and recording sites, rather than as an additional statistical test of individual contact-level effects. This analysis was performed using custom MATLAB code (see “Data and code availability” section).

##### Comparison of decoding accuracies

We compared decoding accuracy based on PAC with decoding based on theta and gamma activity alone. Theta and gamma decoding analyses were performed using the same SEEG contacts included in the spatially clustered regions showing significant PAC decoding, separately for each condition (AL, Phase 1, and Phase 2; Figure 4). For each selected contact and condition, decoding accuracy was obtained separately for PAC, theta, and gamma features using the same decoding procedure described above. For each observation, decoding accuracy was normalized against its corresponding null distribution obtained from 1,000 permutations (z-score). These scores quantify decoding performance relative to the feature-specific null distribution, in units of its standard deviation, and thus enable comparison of standardized decoding evidence across features rather than of raw accuracy on a common absolute scale. Note that for theta and gamma activity, classifiers were trained on time–frequency maps restricted to the corresponding frequency bands and resized to match the dimensions of the PAC comodulograms, thereby matching classifier input dimensions across features. Because gamma spans a broad frequency range (40-150 Hz), we performed separate classifications for low gamma (40-90 Hz) and high gamma (90-150 Hz). Low- and high-gamma decoding accuracies were correlated across contacts in our data (r = 0.52, p < .05), we thus averaged their normalized decoding scores to obtain a composite gamma measure. To evaluate whether decoding performance differed between features while accounting for the non-independence of contacts within participants, we fitted a generalized estimating equation (GEE) with participants as clusters, an exchangeable working correlation structure, and robust sandwich standard errors (R package geepack), with feature (PAC, theta, gamma) and condition (AL, Phase 1, Phase 2) and their interaction as factors. Omnibus effects were evaluated using sequential Wald chi-square tests. Pairwise comparisons between features were performed using custom contrast vectors applied to the model coefficients, with standard errors derived from the robust sandwich covariance matrix and p-values obtained from z-tests. Because only 14 participants contributed, we additionally applied the Kauermann-Carroll (KC) small-sample correction to the sandwich covariance matrix, re-evaluating all contrasts against a t distribution with degrees of freedom equal to the number of clusters minus the number of model parameters (df = 5). Omnibus feature and interaction effects under KC correction were evaluated as F statistics referred to F(df_numerator, 5). All analyses were performed in R. Statistical significance was assessed at α = 0.05.

To test whether PAC decoding after learning exceeded decoding before learning, we compared physically identical stimuli presented before and after the learning phase: sounds during passive exposure (pre-learning) were compared with sounds during Phase 1 (post-learning), and letters during passive exposure (pre-learning) were compared with letters during Phase 2 (post-learning). The associative learning condition was excluded from this comparison as it contains both cues on every trial. Analyses included contacts showing above-chance PAC decoding after learning (visual: 14 contacts; auditory: 24 contacts; 14 participants). Decoding z-scores were analysed using the same GEE framework described above, with participant as the clustering unit, a Gaussian identity link, and an exchangeable working correlation structure. The model included modality (visual, auditory; effect-coded ±0.5) and learning (POST vs PRE; effect-coded ±0.5) and their interaction as fixed effects. Cluster-robust sandwich standard errors were used for inference with z-tests, and the Kauermann-Carroll small-sample correction was applied, re-evaluating all effects against a t distribution with 10 degrees of freedom (clusters minus parameters). All tests were two-sided.

##### Relationship between PAC and Behavior

To test whether PAC-based decoding was behaviorally relevant, normalized decoding accuracy was averaged across significant contacts within each participant and correlated with behavioral performance separately for Phase 1 and Phase 2, using a one-tailed Pearson correlation based on the directional hypothesis that stronger decoding would be associated with better associative memory performance.

## Acknowledgements

This work was supported by a grant from Fonds de Recherche du Québec - Santé and Brain Canada Future Leaders to P.A., and NSERC Discovery grant to P.A.

## Authors contribution

Conceptualization, A.B. and P.A.; Methodology, A.B. and P.A.; Software, A.B., C.L. and P.A.; Formal Analysis, A.B. and P.A.; Investigation, A.B., L.M., P.L.B., P.A.; Resources, P.A., L.M., P.L.B.; Data Curation, A.B.; Writing - Original Draft, A.B.; Writing - Review and Editing, A.B., L.M., P.L-B., C.L. and P.A.; Visualization, A.B.; Supervision, P.A.; Funding Acquisition, P.A.

## Declaration of interest

The authors declare no competing financial or non-financial interests related to the content of this research.

## Inclusion and diversity

We are committed to fostering inclusive, diverse, and equitable practices in all aspects of our research, ensuring that our findings contribute to a broader understanding of human neuroscience.

## Declaration of generative AI and AI-assisted technologies in the writing process

During the preparation of this work, the lead author used ChatGPT 4.o and Claude Opus 5.6 for language, grammar and syntax purposes exclusively. After using this tool, the lead author reviewed and edited the content as need and takes full responsibility for the content of the publication.

## Notes

https://osf.io/d8cuy

### Competing Interest Statement

The authors have declared no competing interest.

### Summary of Updates

The major changes include: 1) Physiological basis of the PAC analysis: We added time-frequency analyses and power spectra demonstrating task-related theta- and gamma-band activity. These analyses provide an empirical basis for the frequency ranges used in the PAC analyses. We also removed stimulus-locked evoked responses prior to the time-frequency and PAC analyses to reduce potential contributions from evoked activity. 2) Validation and interpretation of PAC: We now directly compare PAC-based decoding with decoding based on theta and gamma activity alone. PAC decoding was significantly higher than decoding based on each individual frequency component, suggesting that the information captured by the PAC patterns is not reducible to either component in isolation. 3) Statistical and machine-learning framework: We clarified the statistical level of each analysis, the spatial clustering procedure, and the treatment of dependencies across contacts. For the decoding analyses, we implemented a held-out test set with cross-validation for hyperparameter selection, 1,000 label-shuffling permutations, and additional analyses of classifier stability and feature-to-trial ratios. For group-level inference we now use generalized estimating equations (GEE) with participant as the clustering unit, exchangeable working correlation, and cluster-robust sandwich standard errors, with Kauermann-Carroll small-sample correction applied throughout given the limited number of participants. 4) Relationship between PAC decoding and learning: We now consider the pre-learning sound- and picture-exposure conditions as cue-only controls (before learning), allowing us to distinguish information emerging after associative learning from information related to the sensory cues themselves. 5) Experimental design and methodology: We substantially expanded the description of the tasks structure, trial numbers, stimulus presentation, response procedures, PAC computation, filtering, frequency extraction, normalization, and spatial inference procedures. 6) Conceptual framework and discussion: We clarified how our findings relate to theta-gamma models of sequential coding, cross-frequency binding, and hippocampal-cortical theories of memory, while also refining our interpretation of PAC as a potential representational signature.

