## Supplemental Table 1 to 8 for "Cortico-Hippocampal phase–amplitude coupling is a signature of learned audiovisual associations in humans"

**The PDF file includes:**

Table S1 to S8

Table S1: regions and MNI coordinates Figure 2A.

| **X** | **Y** | **Z** | **HCP** | **AAL3** |
| --- | --- | --- | --- | --- |
| 49 | -19 | 3 | White_Matter | Temporal_Sup_R |
| 56 | -18 | 2 | White_Matter | Temporal_Sup_R |
| 63 | -14 | 4 | White_Matter | Temporal_Sup_R |
| 66 | -13 | 4 | R_A4 | Temporal_Sup_R |
| 51 | -20 | -2 | White_Matter | Temporal_Sup_R |
| 62 | -19 | 1 | R_A5 | Temporal_Sup_R |
| 53 | -7 | -15 | R_STSda | Temporal_Sup_R |
| 58 | -6 | -10 | R_STSda | Temporal_Sup_R |
| 61 | -5 | -9 | R_A5 | Temporal_Sup_R |
| -50 | -13 | -6 | L_STSda | Temporal_Sup_L |
| -53 | -11 | -2 | White_Matter | Temporal_Sup_L |
| -57 | -9 | -1 | L_A5 | Temporal_Sup_L |
| -61 | -6 | 0 | L_A4 | Temporal_Sup_L |
| -62 | -28 | 6 | White_Matter | Temporal_Sup_L |
| 49 | -3 | -13 | White_Matter | Temporal_Sup_R |
| 55 | 0 | -11 | R_STSda | Temporal_Sup_R |
| 60 | 2 | -10 | R_A5 | Temporal_Sup_R |
| -50 | -7 | -4 | L_TA2 | Temporal_Sup_L |
| -56 | -5 | -4 | L_A5 | Temporal_Sup_L |
| -61 | -2 | -5 | L_A5 | Temporal_Sup_L |
| -52 | -1 | -14 | L_STSda | Temporal_Sup_L |
| -57 | -1 | -12 | L_STSda | Temporal_Sup_L |
| -42 | 2 | -11 | L_PoI2 | Temporal_Sup_L |
| 47 | -3 | -11 | R_PI | Temporal_Sup_R |
| 53 | -1 | -10 | White_Matter | Temporal_Sup_R |
| 64 | 1 | -6 | R_A5 | Temporal_Sup_R |
| 44 | -24 | 2 | White_Matter | Temporal_Sup_R |
| 49 | -26 | 5 | White_Matter | Temporal_Sup_R |
| 55 | -28 | 8 | White_Matter | Temporal_Sup_R |
| 62 | -30 | 8 | White_Matter | Temporal_Sup_R |
| 48 | -9 | -11 | R_STSda | Temporal_Sup_R |
| 59 | -7 | -7 | R_A5 | Temporal_Sup_R |
| 64 | -7 | -6 | R_A5 | Temporal_Sup_R |
| 47 | -22 | 2 | White_Matter | Temporal_Sup_R |
| 53 | -20 | 3 | White_Matter | Temporal_Sup_R |
| 59 | -19 | 4 | White_Matter | Temporal_Sup_R |
| 65 | -18 | 4 | R_A5 | Temporal_Sup_R |
| 48 | -10 | -10 | R_STSda | Temporal_Sup_R |
| 55 | -8 | -8 | R_STSda | Temporal_Sup_R |
| 61 | -6 | -4 | R_A5 | Temporal_Sup_R |
| 68 | -5 | -2 | R_A4 | Temporal_Sup_R |
| 56 | -14 | 23 | White_Matter | SupraMarginal_R |
| 38 | -27 | 4 | White_Matter | White_Matter |
| 44 | -27 | 5 | White_Matter | Temporal_Sup_R |
| 51 | -25 | 6 | White_Matter | Temporal_Sup_R |
| -48 | -28 | 5 | L_PBelt | Temporal_Sup_L |
| -56 | -29 | 5 | White_Matter | Temporal_Sup_L |
| 47 | -12 | -3 | R_52 | Temporal_Sup_R |
| 52 | -13 | 0 | White_Matter | Temporal_Sup_R |
| 59 | -14 | 2 | R_PBelt | Temporal_Sup_R |
| 65 | -13 | 4 | R_A4 | Temporal_Sup_R |
| 45 | -31 | 8 | White_Matter | Temporal_Sup_R |
| 52 | -32 | 7 | White_Matter | Temporal_Sup_R |
| 59 | -33 | 8 | R_A5 | Temporal_Sup_R |
| 66 | -33 | 8 | R_A4 | Temporal_Sup_R |
| 51 | -16 | -5 | R_STSda | Temporal_Sup_R |
| 58 | -17 | -5 | R_A5 | Temporal_Sup_R |
| 63 | -17 | -2 | R_A5 | Temporal_Sup_R |
| 70 | -17 | 0 | R_A5 | Temporal_Sup_R |
| 54 | -6 | 0 | R_TA2 | Temporal_Sup_R |
| 60 | -6 | 1 | R_A4 | Temporal_Sup_R |
| 67 | -7 | 3 | R_A4 | Temporal_Sup_R |
| 63 | -37 | 8 | R_TPOJ1 | Temporal_Sup_R |
| 68 | -35 | 13 | R_A4 | Temporal_Sup_R |
| 71 | -4 | -13 | White_Matter | Temporal_Sup_R |
| 60 | -47 | 20 | R_STV | Temporal_Sup_R |
| 46 | -16 | -5 | White_Matter | Temporal_Sup_R |
| 46 | -20 | -1 | White_Matter | Temporal_Sup_R |
| 51 | -36 | 21 | White_Matter | Temporal_Sup_R |
| 58 | -36 | 21 | White_Matter | Temporal_Sup_R |
| 65 | -37 | 21 | R_PSL | Temporal_Sup_R |
| 68 | -38 | 21 | R_PSL | Temporal_Sup_R |
| 44 | -21 | 4 | White_Matter | Temporal_Sup_R |
| 51 | -20 | 4 | R_A1 | Temporal_Sup_R |
| 57 | -19 | 4 | White_Matter | Temporal_Sup_R |
| 64 | -18 | 5 | White_Matter | Temporal_Sup_R |
| 68 | -18 | 6 | R_A4 | Temporal_Sup_R |

Table S2: regions and MNI coordinates Figure 2B

| **X** | **Y** | **Z** | **HCP** | **AAL3** |
| --- | --- | --- | --- | --- |
| 42 | -64 | 28 | R_PGi | Occipital_Mid_R |
| 47 | -67 | 26 | R_PGi | Occipital_Mid_R |
| 11 | -78 | 3 | White_Matter | Calcarine_R |
| 21 | -77 | -1 | White_Matter | Lingual_R |
| 38 | -67 | 11 | White_Matter | Occipital_Mid_R |

Table S3: regions and MNI coordinates Figure 2C

| **X** | **Y** | **Z** | **HCP** | **AAL3** |
| --- | --- | --- | --- | --- |
| 40 | -25 | -15 | White_Matter | Hippocampus_R |
| -28 | -14 | -23 | L_H | Hippocampus_L |
| 20 | -6 | -20 | White_Matter | Hippocampus_R |
| 33 | -13 | -21 | R_H | Hippocampus_R |
| -25 | -13 | -23 | L_H | Hippocampus_L |
| 22 | -26 | -12 | R_H | Hippocampus_R |
| 33 | -25 | -9 | White_Matter | Hippocampus_R |
| -31 | -19 | -17 | White_Matter | Hippocampus_L |
| -23 | -36 | -5 | L_H | Hippocampus_L |
| -33 | -34 | -8 | L_H | Hippocampus_L |
| 26 | -4 | -26 | R_H | Hippocampus_R |
| -17 | -8 | -20 | White_Matter | Hippocampus_L |
| -28 | -5 | -21 | L_H | Hippocampus_L |
| 36 | -19 | -18 | R_H | Hippocampus_R |
| -31 | -19 | -18 | L_H | Hippocampus_L |
| 27 | -33 | -9 | R_H | Hippocampus_R |
| 38 | -30 | -8 | White_Matter | Hippocampus_R |
| -30 | -29 | -12 | L_H | Hippocampus_L |
| -31 | -31 | -12 | L_H | Hippocampus_L |
| -33 | -37 | -3 | White_Matter | Hippocampus_L |
| -27 | -16 | -19 | White_Matter | Hippocampus_L |
| -37 | -16 | -19 | L_H | Hippocampus_L |
| 21 | -3 | -23 | White_Matter | Hippocampus_R |
| 34 | -16 | -23 | R_H | Hippocampus_R |
| 29 | -32 | -7 | White_Matter | Hippocampus_R |
| 30 | -5 | -26 | White_Matter | Hippocampus_R |
| 39 | -11 | -15 | White_Matter | Hippocampus_R |
| -23 | -13 | -18 | White_Matter | Hippocampus_L |
| -35 | -13 | -20 | L_H | Hippocampus_L |
| 23 | -5 | -20 | R_H | Hippocampus_R |
| 33 | -5 | -21 | White_Matter | Hippocampus_R |
| -20 | -6 | -19 | White_Matter | Hippocampus_L |
| 27 | -15 | -20 | White_Matter | Hippocampus_R |
| 35 | -15 | -19 | R_H | Hippocampus_R |
| -34 | -20 | -19 | L_H | Hippocampus_L |
| 31 | -31 | -10 | R_H | Hippocampus_R |
| 40 | -31 | -11 | White_Matter | Hippocampus_R |
| 24 | -27 | -11 | R_H | Hippocampus_R |
| 35 | -30 | -10 | R_H | Hippocampus_R |
| -21 | -15 | -17 | White_Matter | Hippocampus_L |
| -31 | -13 | -16 | White_Matter | Hippocampus_L |
| -25 | -18 | -19 | L_H | Hippocampus_L |
| -35 | -21 | -18 | L_H | Hippocampus_L |
| 33 | -15 | -21 | R_H | Hippocampus_R |
| -25 | -17 | -20 | L_H | Hippocampus_L |
| -36 | -17 | -17 | White_Matter | Hippocampus_L |
| 27 | -19 | -18 | R_H | Hippocampus_R |
| 39 | -19 | -18 | R_H | Hippocampus_R |
| 36 | -35 | -7 | White_Matter | Hippocampus_R |
| 32 | -6 | -26 | White_Matter | Hippocampus_R |
| -27 | -21 | -12 | White_Matter | Hippocampus_L |

Table S4: regions and MNI coordinates Figure 2D

| **X** | **Y** | **Z** | **HCP** | **AAL3** |
| --- | --- | --- | --- | --- |
| 36 | 13 | -10 | R_AAIC | Insula_R |
| 41 | 4 | -10 | R_PoI2 | Insula_R |
| 38 | 9 | -5 | R_PoI2 | Insula_R |
| 20 | -6 | -20 | White_Matter | Hippocampus_R |
| 30 | -3 | -22 | R_H | Amygdala_R |
| 39 | 0 | -22 | White_Matter | White_Matter |
| -44 | -9 | -22 | White_Matter | White_Matter |
| -39 | 8 | -11 | L_AAIC | Insula_L |
| 62 | -19 | 1 | R_A5 | Temporal_Sup_R |
| -45 | -31 | -13 | White_Matter | White_Matter |
| 36 | -26 | -27 | R_TF | Fusiform_R |
| 44 | -25 | -26 | R_TE2p | Fusiform_R |
| 51 | -22 | -23 | White_Matter | Temporal_Inf_R |
| 62 | -20 | -23 | R_TE1m | Temporal_Inf_R |
| 46 | -9 | -19 | White_Matter | White_Matter |
| 53 | -7 | -15 | R_STSda | Temporal_Sup_R |
| 58 | -6 | -10 | R_STSda | Temporal_Sup_R |
| 35 | -55 | -12 | White_Matter | Fusiform_R |
| 39 | -6 | 8 | R_PoI2 | Insula_R |
| 48 | -5 | 10 | R_43 | Rolandic_Oper_R |
| -51 | -12 | -16 | L_STSva | Temporal_Mid_L |
| -61 | -15 | -14 | L_TE1a | Temporal_Mid_L |
| -29 | 13 | 2 | White_Matter | White_Matter |
| -23 | -36 | -5 | L_H | Hippocampus_L |
| -63 | -28 | -14 | White_Matter | Temporal_Mid_L |
| -15 | 50 | 31 | L_9p | Frontal_Sup_2_L |
| -53 | -11 | -2 | White_Matter | Temporal_Sup_L |
| -57 | -9 | -1 | L_A5 | Temporal_Sup_L |
| 35 | 7 | -6 | White_Matter | White_Matter |
| 37 | -4 | -11 | R_PoI1 | White_Matter |
| 34 | -14 | 9 | White_Matter | Insula_R |
| -56 | -5 | -4 | L_A5 | Temporal_Sup_L |
| -17 | -8 | -20 | White_Matter | Hippocampus_L |
| -45 | -2 | -17 | L_PI | Temporal_Mid_L |
| -11 | 48 | 34 | White_Matter | Frontal_Sup_2_L |
| -57 | -1 | -12 | L_STSda | Temporal_Sup_L |
| -34 | 10 | -37 | L_TGd | Temporal_Inf_L |
| -60 | -36 | 5 | L_A5 | Temporal_Mid_L |
| -65 | -36 | 5 | L_A5 | Temporal_Mid_L |
| -13 | 26 | 59 | L_SFL | Frontal_Sup_2_L |
| -6 | 30 | -23 | White_Matter | Rectus_L |
| -60 | 4 | 13 | White_Matter | Postcentral_L |
| -62 | 5 | 15 | L_43 | Postcentral_L |
| -46 | -15 | -18 | White_Matter | White_Matter |
| -55 | -13 | -15 | L_STSva | Temporal_Mid_L |
| -41 | 1 | -36 | L_TGv | Temporal_Inf_L |
| -44 | 17 | 13 | White_Matter | Frontal_Inf_Oper_L |
| 21 | -3 | -23 | White_Matter | Hippocampus_R |
| 65 | -14 | -22 | R_TE1a | Temporal_Mid_R |
| 65 | -25 | -18 | R_TE1m | Temporal_Mid_R |
| -36 | 14 | -5 | White_Matter | Insula_L |
| 53 | -20 | 3 | White_Matter | Temporal_Sup_R |
| 55 | 24 | 20 | R_45 | Frontal_Inf_Tri_R |
| -53 | 1 | -21 | L_STSda | Temporal_Mid_L |
| 62 | -27 | -11 | White_Matter | Temporal_Mid_R |
| 32 | -9 | 7 | White_Matter | Putamen_R |
| -52 | -3 | 17 | White_Matter | Precentral_L |
| -60 | -2 | 20 | L_4 | Postcentral_L |
| 51 | 29 | 9 | R_45 | Frontal_Inf_Tri_R |
| 58 | 28 | 9 | R_45 | Frontal_Inf_Tri_R |
| 55 | 25 | 14 | White_Matter | Frontal_Inf_Tri_R |
| -42 | -13 | -16 | White_Matter | White_Matter |
| -21 | -15 | -17 | White_Matter | Hippocampus_L |
| 38 | 38 | 21 | White_Matter | Frontal_Mid_2_R |
| -43 | -49 | 29 | White_Matter | Angular_L |
| -45 | -6 | -29 | White_Matter | Temporal_Inf_L |
| -23 | -37 | -9 | White_Matter | ParaHippocampal_L |
| -30 | 18 | 13 | L_FOP4 | Insula_L |
| -14 | 39 | 33 | White_Matter | Frontal_Sup_2_L |
| -41 | 5 | -38 | L_TGd | Temporal_Inf_L |
| -12 | 26 | 52 | L_8BL | Frontal_Sup_2_L |
| -21 | 30 | 62 | White_Matter | Frontal_Sup_2_L |
| -7 | 30 | -24 | L_OFC | Rectus_L |
| -38 | 18 | 12 | L_FOP4 | Frontal_Inf_Tri_L |
| 44 | 37 | 18 | R_p9-46v | Frontal_Mid_2_R |
| -62 | -11 | -20 | L_TE1a | Temporal_Mid_L |
| -59 | -29 | -16 | White_Matter | Temporal_Mid_L |
| -69 | -29 | 8 | L_A5 | Temporal_Mid_L |
| 47 | 38 | 19 | R_p9-46v | Frontal_Mid_2_R |
| 66 | -30 | -8 | R_TE1p | Temporal_Mid_R |
| 37 | -51 | -12 | White_Matter | Fusiform_R |
| 46 | -3 | 16 | R_FOP2 | Rolandic_Oper_R |
| -38 | 2 | -12 | L_Pir | Insula_L |
| -35 | 6 | 0 | White_Matter | White_Matter |
| 51 | -16 | -5 | R_STSda | Temporal_Sup_R |
| -47 | -31 | -16 | White_Matter | Temporal_Mid_L |
| 47 | -14 | -23 | White_Matter | White_Matter |
| 59 | -11 | -22 | R_TE1a | Temporal_Mid_R |
| 44 | -26 | -25 | R_TE2p | Fusiform_R |
| 51 | -25 | -25 | White_Matter | Temporal_Inf_R |
| 58 | -24 | -25 | R_TE2a | Temporal_Inf_R |
| 30 | -11 | 3 | White_Matter | Putamen_R |
| 28 | -12 | 10 | White_Matter | Putamen_R |
| -66 | -34 | -10 | L_TE1p | Temporal_Mid_L |
| -52 | -52 | 24 | White_Matter | SupraMarginal_L |

Table S5: regions and MNI coordinates Figure 2E

| **X** | **Y** | **Z** | **HCP** | **AAL3** |
| --- | --- | --- | --- | --- |
| 30 | -3 | -22 | R_H | Amygdala_R |
| 62 | -19 | 1 | R_A5 | Temporal_Sup_R |
| -37 | -32 | -13 | L_PHA2 | Temporal_Inf_L |
| -45 | -31 | -13 | White_Matter | White_Matter |
| -53 | -28 | -11 | White_Matter | Temporal_Mid_L |
| -30 | -29 | -12 | L_H | Hippocampus_L |
| 39 | 1 | -21 | R_PoI1 | White_Matter |
| 65 | -18 | 4 | R_A5 | Temporal_Sup_R |
| 51 | 35 | 0 | R_IFSa | Frontal_Inf_Tri_R |
| 58 | 28 | 9 | R_45 | Frontal_Inf_Tri_R |
| 47 | -27 | 24 | R_PFcm | SupraMarginal_R |
| 52 | 30 | 4 | White_Matter | Frontal_Inf_Tri_R |
| -32 | 19 | 10 | L_FOP4 | Frontal_Inf_Tri_L |
| -38 | 18 | 12 | L_FOP4 | Frontal_Inf_Tri_L |
| -31 | 13 | 8 | L_MI | Insula_L |
| 33 | 3 | -16 | White_Matter | White_Matter |
| 59 | -14 | 2 | R_PBelt | Temporal_Sup_R |
| 28 | -56 | 61 | White_Matter | Parietal_Sup_R |
| 35 | -5 | 3 | White_Matter | Putamen_R |
| 34 | -9 | 10 | R_Ig | Insula_R |
| 55 | -26 | 25 | R_PF | SupraMarginal_R |
| 62 | -24 | 25 | R_PFop | SupraMarginal_R |
| 19 | -62 | 50 | White_Matter | Parietal_Sup_R |
| 24 | -58 | 56 | R_LIPv | Parietal_Sup_R |
| 35 | -1 | -6 | White_Matter | White_Matter |
| 34 | -4 | 2 | White_Matter | Putamen_R |

Table S6: regions and coordinates Figure 4A

| **X** | **Y** | **Z** | **HCP** | **AAL3** |
| --- | --- | --- | --- | --- |
| -35 | -10 | -22 | White_Matter | White_Matter |
| 36 | -19 | -18 | R_H | Hippocampus_R |
| 34 | -16 | -23 | R_H | Hippocampus_R |
| 32 | 15 | 6 | R_MI | Insula_R |
| -38 | -18 | -23 | White_Matter | Fusiform_L |
| 34 | 17 | 2 | R_MI | Insula_R |

Table S7: regions and MNI coordinates Figure 4B

| **X** | **Y** | **Z** | **HCP** | **AAL3** |
| --- | --- | --- | --- | --- |
| 40 | -39 | -14 | White_Matter | White_Matter |
| 56 | -18 | 2 | White_Matter | Temporal_Sup_R |
| -25 | -35 | -13 | White_Matter | ParaHippocampal_L |
| 46 | -9 | -19 | White_Matter | White_Matter |
| -61 | -15 | -14 | L_TE1a | Temporal_Mid_L |
| -63 | -28 | -14 | White_Matter | Temporal_Mid_L |
| -30 | -29 | -12 | L_H | Hippocampus_L |
| 40 | -7 | -7 | R_PoI2 | Insula_R |
| -49 | 39 | -4 | White_Matter | Frontal_Inf_Orb_2_L |
| 44 | -5 | -23 | White_Matter | White_Matter |
| 36 | -2 | -3 | White_Matter | Putamen_R |
| 31 | -12 | 13 | White_Matter | White_Matter |
| -49 | 38 | -5 | White_Matter | Frontal_Inf_Orb_2_L |
| -62 | -11 | -20 | L_TE1a | Temporal_Mid_L |
| -63 | -26 | -13 | White_Matter | Temporal_Mid_L |
| 65 | -13 | 4 | R_A4 | Temporal_Sup_R |
| 66 | -33 | 8 | R_A4 | Temporal_Sup_R |
| 46 | -39 | -11 | White_Matter | White_Matter |
| 58 | -17 | -5 | R_A5 | Temporal_Sup_R |
| 70 | -17 | 0 | R_A5 | Temporal_Sup_R |
| 55 | -17 | -11 | R_STSva | Temporal_Mid_R |
| 63 | -37 | 8 | R_TPOJ1 | Temporal_Sup_R |
| 51 | -20 | 4 | R_A1 | Temporal_Sup_R |
| 31 | -7 | 15 | White_Matter | White_Matter |

Table S8: regions and coordinates Figure 4C

| **X** | **Y** | **Z** | **HCP** | **AAL3** |
| --- | --- | --- | --- | --- |
| 18 | -38 | 27 | White_Matter | White_Matter |
| -39 | 8 | -11 | L_AAIC | Insula_L |
| -35 | 5 | -11 | White_Matter | White_Matter |
| -33 | 8 | -4 | White_Matter | White_Matter |
| 18 | -43 | 28 | White_Matter | White_Matter |
| 7 | -25 | 36 | White_Matter | Cingulate_Mid_R |
| 52 | 28 | 11 | R_45 | Frontal_Inf_Tri_R |
| 64 | -28 | -10 | R_TE1m | Temporal_Mid_R |
| 58 | 28 | 9 | R_45 | Frontal_Inf_Tri_R |
| 7 | -26 | 41 | R_23c | Cingulate_Mid_R |
| 35 | -30 | -10 | R_H | Hippocampus_R |
| 66 | -30 | -8 | R_TE1p | Temporal_Mid_R |
| 36 | -35 | -7 | White_Matter | Hippocampus_R |
| -35 | 6 | 0 | White_Matter | White_Matter |
